## Supplementary Materials for "Specific Modulation of CRISPR Transcriptional Activators through RNA-Sensing Guide RNAs in Mammalian Cells and Zebrafish Embryos"

### Relevant RNA sequences and primers

#### 1. Native sgRNA sequences

+G Spacer scaffold

sgRNA 1:
GCAGTCGCGTGTAGCGAAGCAGTTTTAGAGCTAGAAATAGCAAGTTAAAATAAGGCTAGTCC
GTTATCAACTTGAAAAAGTGGCACCGAGTCGGTGC
sgRNA 2:
GTAAGTCGGAGTACTGTCCTGTTTTAGAGCTAGAAATAGCAAGTTAAAATAAGGCTAGTCCG
TTATCAACTTGAAAAAGTGGCACCGAGTCGGTGC
sgRNA 3:
GCATGACATTATTCCCGCCTCGTTTTAGAGCTAGAAATAGCAAGTTAAAATAAGGCTAGTCC
GTTATCAACTTGAAAAAGTGGCACCGAGTCGGTGC
sgRNA 4:
GCCAGGTAATGACTAGCAGGAGTTTTAGAGCTAGAAATAGCAAGTTAAAATAAGGCTAGTCC
GTTATCAACTTGAAAAAGTGGCACCGAGTCGGTGC
sgRNA 5:
GCACTACTTCAGTAAGAGTCGTTTTAGAGCTAGAAATAGCAAGTTAAAATAAGGCTAGTCCG
TTATCAACTTGAAAAAGTGGCACCGAGTCGGTGC

*For sgRNAs with spacer sequences starting with an A, C or T, an extra G was added for facilitating expression from U6 vectors

#### 2. Native sgRNAs with randomised spacer sequences

+G randomised nt Spacer scaffold

sgRNA 1 (D1):
GAGTCGCGTGTAGCGAAGCAGTTTTAGAGCTAGAAATAGCAAGTTAAAATAAGGCTAGTCCG
TTATCAACTTGAAAAAGTGGCACCGAGTCGGTGC

sgRNA 1 (D2):
GTGGTCGCGTGTAGCGAAGCAGTTTTAGAGCTAGAAATAGCAAGTTAAAATAAGGCTAGTCC
GTTATCAACTTGAAAAAGTGGCACCGAGTCGGTGC

sgRNA 1 (D3):
GTGATCGCGTGTAGCGAAGCAGTTTTAGAGCTAGAAATAGCAAGTTAAAATAAGGCTAGTCC
GTTATCAACTTGAAAAAGTGGCACCGAGTCGGTGC

sgRNA 2 (D1):
GCTAAGTCGGAGTACTGTCCTGTTTTAGAGCTAGAAATAGCAAGTTAAAATAAGGCTAGTCC
GTTATCAACTTGAAAAAGTGGCACCGAGTCGGTGC

sgRNA 2 (D2):
GCAAAGTCGGAGTACTGTCCTGTTTTAGAGCTAGAAATAGCAAGTTAAAATAAGGCTAGTCC
GTTATCAACTTGAAAAAGTGGCACCGAGTCGGTGC

sgRNA 2 (D3):
GCATAGTCGGAGTACTGTCCTGTTTTAGAGCTAGAAATAGCAAGTTAAAATAAGGCTAGTCC
GTTATCAACTTGAAAAAGTGGCACCGAGTCGGTGC

sgRNA 3 (D1):
GATGACATTATTCCCGCCTCGTTTTAGAGCTAGAAATAGCAAGTTAAAATAAGGCTAGTCCG
TTATCAACTTGAAAAAGTGGCACCGAGTCGGTGC

sgRNA 3 (D2):
GGTGACATTATTCCCGCCTCGTTTTAGAGCTAGAAATAGCAAGTTAAAATAAGGCTAGTCCG
TTATCAACTTGAAAAAGTGGCACCGAGTCGGTGC

sgRNA 3 (D3):
GGGGACATTATTCCCGCCTCGTTTTAGAGCTAGAAATAGCAAGTTAAAATAAGGCTAGTCCG
TTATCAACTTGAAAAAGTGGCACCGAGTCGGTGC

#### 3. First-generation iSBH-sgRNA sequences

+GC spacer* loop spacer scaffold

*An additional GC sequence was added to all iSBH-sgRNA sequences. G helps expression from U6 promoter, while C base-pairs with the first C from the scaffold

First-generation iSBH-sgRNA 1:
GCTGCTTCGCTAGTCGCTTCTGCACACTACGCATCCCAGTCGCGTGTAGCGAAGCAGTTTTA
GAGCTAGAAATAGCAAGTTAAAATAAGGCTAGTCCGTTATCAACTTGAAAAAGTGGCACCGA
GTCGGTGC

First-generation iSBH-sgRNA 2:
GCAGGACAGTACAGCGAGATACGAACATCCCTAACAGTAAGTCGGAGTACTGTCCTGTTTTA
GAGCTAGAAATAGCAAGTTAAAATAAGGCTAGTCCGTTATCAACTTGAAAAAGTGGCACCGA
GTCGGTGC

First-generation iSBH-sgRNA 3:
GCGAGGCGGGAAGTATGGAATGGACCAGTGAATACACATGACATTATTCCCGCCTCGTTTTA
GAGCTAGAAATAGCAAGTTAAAATAAGGCTAGTCCGTTATCAACTTGAAAAAGTGGCACCGA
GTCGGTGC

First-generation iSBH-sgRNA 4:
GCTCCTGCTAGTACTTAAATGGGCTCCCCACATCAACCAGGTAATGACTAGCAGGAGTTTTA
GAGCTAGAAATAGCAAGTTAAAATAAGGCTAGTCCGTTATCAACTTGAAAAAGTGGCACCGA
GTCGGTGC

First-generation iSBH-sgRNA 5:
GCGACTCTTACTCCAGTTCTGCATCTTGTTCGTCCAGCACTACTTCAGTAAGAGTCGTTTTA
GAGCTAGAAATAGCAAGTTAAAATAAGGCTAGTCCGTTATCAACTTGAAAAAGTGGCACCGA
GTCGGTGC

#### 4. Second-generation iSBH-sgRNA sequences

+GC spacer* extension*loop extensionspacer scaffold

Backfold=spacer*+extension+loop

*An additional GC sequence was added to all iSBH-sgRNA sequences. G helps expression from U6 promoter, while C base-pairs with the first C from the scaffold

Second-generation iSBH-sgRNA 1:
GCTGCTTCGCTAGTCGCTTCTGTCCGTACGTTCACACTACGCATCCAACTGACGTTCAGTCG
CGTGTAGCGAAGCAGTTTTAGAGCTAGAAATAGCAAGTTAAAATAAGGCTAGTCCGTTATCA
ACTTGAAAAAGTGGCACCGAGTCGGTGC

Second-generation iSBH-sgRNA 2:
GCAGGACAGTACAGCGAGATACCAGCTTAGTCGAACATCCCTAACAGACATAGCCCGTAAGT
CGGAGTACTGTCCTGTTTTAGAGCTAGAAATAGCAAGTTAAAATAAGGCTAGTCCGTTATCA
ACTTGAAAAAGTGGCACCGAGTCGGTGC

Second-generation iSBH-sgRNA 3:
GCGAGGCGGGAAGTATGGAATGAGTGGGATGGGACCAGTGAATACACCAAACCAGACATGAC
ATTATTCCCGCCTCGTTTTAGAGCTAGAAATAGCAAGTTAAAATAAGGCTAGTCCGTTATCA
ACTTGAAAAAGTGGCACCGAGTCGGTGC

#### 5. Modular iSBH-sgRNA sequences

+GC spacer* trigger-sensing region extension spacer scaffold

*An additional GC sequence was added to all iSBH-sgRNA sequences. G helps expression from U6 promoter, while C base-pairs with the first C from the scaffold

Modular iSBH-sgRNA (trigger D/sgRNA2/loop=14nt):
GCAGGACAGTACAGCGAGACAGTGTTCTCTGCACAGATAAGGACAAGCAATGAAATCTGAGT
CGGAGTACTGTCCTGTTTTAGAGCTAGAAATAGCAAGTTAAAATAAGGCTAGTCCGTTATCA
ACTTGAAAAAGTGGCACCGAGTCGGTGC

Modular iSBH-sgRNA (trigger D/sgRNA3/loop=14nt):
GCGAGGCGGGAAGTATGGACAGTGTTCTCTGCACAGATAAGGACAAGCATTGAAATCTGGAC
ATTATTCCCGCCTCGTTTTAGAGCTAGAAATAGCAAGTTAAAATAAGGCTAGTCCGTTATCA
ACTTGAAAAAGTGGCACCGAGTCGGTGC

Modular iSBH-sgRNA (trigger D/sgRNA5/loop=14nt):
GCGACTCTTACTCCAGTCAGTGTTCTCTGCACAGATAAGGACAAACGTGATGAGTTCACCTA
CTTCAGTAAGAGTCGTTTTAGAGCTAGAAATAGCAAGTTAAAATAAGGCTAGTCCGTTATCA
ACTTGAAAAAGTGGCACCGAGTCGGTGC

Modular iSBH-sgRNA (trigger A/sgRNA2/loop=14nt (1)):
GCAGGACAGTACAACGAAAACTGATCATCACCACAACACAACTAAAGGTCTTGAGAAGTAGT
CGGAGTACTGTCCTGTTTTAGAGCTAGAAATAGCAAGTTAAAATAAGGCTAGTCCGTTATCA
ACTTGAAAAAGTGGCACCGAGTCGGTGC

Modular iSBH-sgRNA (trigger A/sgRNA2/loop=14nt (2)):

GCAGGACAGTACAACGAACGACTAAACTGATCATCACCACAACACGGATATGTTGCGTCAGT
CGGAGTACTGTCCTGTTTTAGAGCTAGAAATAGCAAGTTAAAATAAGGCTAGTCCGTTATCA
ACTTGAAAAAGTGGCACCGAGTCGGTGC

Modular iSBH-sgRNA (trigger A/sgRNA2/loop=16nt):
GCAGGACAGTACAGCGAACGACTAAACTGATCATCACCACAACACAACGATATGTTGCGTCA
GTCGGAGTACTGTCCTGTTTTAGAGCTAGAAATAGCAAGTTAAAATAAGGCTAGTCCGTTAT
CAACTTGAAAAAGTGGCACCGAGTCGGTGC

Modular iSBH-sgRNA (trigger A/sgRNA2/loop=18nt):
GCAGGACAGTACAGCGAACGACTAAACTGATCATCACCACAACACAACTAGATATGTTGCGT
CAGTCGGAGTACTGTCCTGTTTTAGAGCTAGAAATAGCAAGTTAAAATAAGGCTAGTCCGTT
ATCAACTTGAAAAAGTGGCACCGAGTCGGTGC

Modular iSBH-sgRNA (trigger A/sgRNA2/loop=20nt):
GCAGGACAGTACAACGAACGACTAAACTGATCATCACCACAACACAACTAAAGATATGTTGC
GTCAGTCGGAGTACTGTCCTGTTTTAGAGCTAGAAATAGCAAGTTAAAATAAGGCTAGTCCG
TTATCAACTTGAAAAAGTGGCACCGAGTCGGTGC

Modular iSBH-sgRNA (trigger A/sgRNA2/loop=22nt):
GCAGGACAGTACAGCGAAACACAACTAAACTGATCATCACCACAACACAACTAACAGCGTAG
GAGTGAGTCGGAGTACTGTCCTGTTTTAGAGCTAGAAATAGCAAGTTAAAATAAGGCTAGTC
CGTTATCAACTTGAAAAAGTGGCACCGAGTCGGTGC

Modular iSBH-sgRNA (trigger A/sgRNA2/loop=24nt):
GCAGGACAGTACAGCGAAACACAACTAAACTGATCATCACCACAACACAACTAAACCAGCGT
AGGAGTGAGTCGGAGTACTGTCCTGTTTTAGAGCTAGAAATAGCAAGTTAAAATAAGGCTAG
TCCGTTATCAACTTGAAAAAGTGGCACCGAGTCGGTGC

Modular iSBH-sgRNA (trigger A/sgRNA2/loop=26nt):
GCAGGACAGTACAGCGAAACACAACTAAACTGATCATCACCACAACACAACTAAACTGCAGC
GTAGGAGTGAGTCGGAGTACTGTCCTGTTTTAGAGCTAGAAATAGCAAGTTAAAATAAGGCT
AGTCCGTTATCAACTTGAAAAAGTGGCACCGAGTCGGTGC

Modular iSBH-sgRNA (trigger A/sgRNA2/loop=28nt):
GCAGGACAGTACAGCGAAAACTGATCATCACCACAACACAACTAAACTGATCATCACCACGG
TCTTGAGAAGTAGTCGGAGTACTGTCCTGTTTTAGAGCTAGAAATAGCAAGTTAAAATAAGG
CTAGTCCGTTATCAACTTGAAAAAGTGGCACCGAGTCGGTGC

Modular iSBH-sgRNA (trigger A/sgRNA2/loop=30nt):
GCAGGACAGTACAGCGAAAACTGATCATCACCACAACACAACTAAACTGATCATCACCACAA
GGTCTTGAGAAGTAGTCGGAGTACTGTCCTGTTTTAGAGCTAGAAATAGCAAGTTAAAATAA
GGCTAGTCCGTTATCAACTTGAAAAAGTGGCACCGAGTCGGTGC

Modular iSBH-sgRNA (trigger A/sgRNA2/loop=32nt):
GCAGGACAGTACAGCGAAAACTGATCATCACCACAACACAACTAAACTGATCATCACCACAA
CAGGTCTTGAGAAGTAGTCGGAGTACTGTCCTGTTTTAGAGCTAGAAATAGCAAGTTAAAAT
AAGGCTAGTCCGTTATCAACTTGAAAAAGTGGCACCGAGTCGGTGC

Modular iSBH-sgRNA (trigger A/sgRNA2/loop=34nt):

GCAGGACAGTACAGCGAAAACTGATCATCACCACAACACAACTAAACTGATCATCACCACAA
CACAGGTCTTGAGAAGTAGTCGGAGTACTGTCCTGTTTTAGAGCTAGAAATAGCAAGTTAAA
ATAAGGCTAGTCCGTTATCAACTTGAAAAAGTGGCACCGAGTCGGTGC

Modular iSBH-sgRNA (trigger B/sgRNA2/loop=14nt (1)):
GCAGGACAGTACGACGAGATGAAGTGAAAAACAGAGGAAGGAAGGAGTTGGTCAAGTCAAGT
CGGAGTACTGTCCTGTTTTAGAGCTAGAAATAGCAAGTTAAAATAAGGCTAGTCCGTTATCA
ACTTGAAAAAGTGGCACCGAGTCGGTGC

Modular iSBH-sgRNA (trigger B/sgRNA2/loop=14nt (2)):
GCAGGACAGTACAACGAACAACGACCAAATGTCTCTAGTCATCCCCACAGGTGGAAGTTAGT
CGGAGTACTGTCCTGTTTTAGAGCTAGAAATAGCAAGTTAAAATAAGGCTAGTCCGTTATCA
ACTTGAAAAAGTGGCACCGAGTCGGTGC

Modular iSBH-sgRNA (trigger B/sgRNA2/loop=16nt):
GCAGGACAGTACAGCGAGGGGCAGATGAAGTGAAAAACAGAGGAAGGACACGGCATAGGCCA
GTCGGAGTACTGTCCTGTTTTAGAGCTAGAAATAGCAAGTTAAAATAAGGCTAGTCCGTTAT
CAACTTGAAAAAGTGGCACCGAGTCGGTGC

Modular iSBH-sgRNA (trigger B/sgRNA2/loop=18nt):
GCAGGACAGTACAGCGAGGGCAGATGAAGTGAAAAACAGAGGAAGGAAGGTCAAGTCAGATG
CAGTCGGAGTACTGTCCTGTTTTAGAGCTAGAAATAGCAAGTTAAAATAAGGCTAGTCCGTT
ATCAACTTGAAAAAGTGGCACCGAGTCGGTGC

Modular iSBH-sgRNA (trigger B/sgRNA2/loop=20nt):
GCAGGACAGTACAGCGAGGGGCAGATGAAGTGAAAAACAGAGGAAGGAAGGACACGGCATAG
GCCAGTCGGAGTACTGTCCTGTTTTAGAGCTAGAAATAGCAAGTTAAAATAAGGCTAGTCCG
TTATCAACTTGAAAAAGTGGCACCGAGTCGGTGC

Modular iSBH-sgRNA (trigger B/sgRNA2/loop=22nt):
GCAGGACAGTACAGCGAGATGAAGTGAAAAACAGAGGAAGGAAGGATTGTGGGAGTTGGTCA
AGTCAAGTCGGAGTACTGTCCTGTTTTAGAGCTAGAAATAGCAAGTTAAAATAAGGCTAGTC
CGTTATCAACTTGAAAAAGTGGCACCGAGTCGGTGC

Modular iSBH-sgRNA (trigger B/sgRNA2/loop=24nt):
GCAGGACAGTACAGCGAGATGGGGGCAGATGAAGTGAAAAACAGAGGAAGGAAGGATCAGAT
GCAACCAAGTCGGAGTACTGTCCTGTTTTAGAGCTAGAAATAGCAAGTTAAAATAAGGCTAG
TCCGTTATCAACTTGAAAAAGTGGCACCGAGTCGGTGC

Modular iSBH-sgRNA (trigger B/sgRNA2/loop=26nt):
GCAGGACAGTACAGCGAGGCGATCACAACGACCAAATGTCTCTAGTCATCCCCTGCTTGTCA
GTGTATTCGAGTCGGAGTACTGTCCTGTTTTAGAGCTAGAAATAGCAAGTTAAAATAAGGCT
AGTCCGTTATCAACTTGAAAAAGTGGCACCGAGTCGGTGC

Modular iSBH-sgRNA (trigger B/sgRNA2/loop=28nt):
GCAGGACAGTACAGCGAGATGAAGTGAAAAACAGAGGAAGGAAGGATTGTGGGATTATGAGT
TGGTCAAGTCAAGTCGGAGTACTGTCCTGTTTTAGAGCTAGAAATAGCAAGTTAAAATAAGG
CTAGTCCGTTATCAACTTGAAAAAGTGGCACCGAGTCGGTGC

Modular iSBH-sgRNA (trigger C/sgRNA2/loop=14nt (1)):

GCAGGACAGTACGACGAAGACAGACTGTGAAAGGAGAGGCGAGGGATTTATCAGGATGTAGT
CGGAGTACTGTCCTGTTTTAGAGCTAGAAATAGCAAGTTAAAATAAGGCTAGTCCGTTATCA
ACTTGAAAAAGTGGCACCGAGTCGGTGC

Modular iSBH-sgRNA (trigger C/sgRNA2/loop=14nt (2)):
GCAGGACAGTACGACGAAGTGCGCAGCGCGCCTCTCTCTCTCGCTCGGCAAGCTAAGCAAGT
CGGAGTACTGTCCTGTTTTAGAGCTAGAAATAGCAAGTTAAAATAAGGCTAGTCCGTTATCA
ACTTGAAAAAGTGGCACCGAGTCGGTGC

Modular iSBH-sgRNA (trigger C/sgRNA2/loop=16nt):
GCAGGACAGTACAGCGAACAGACAGACTGTGAAAGGAGAGGCGAGGGATCAATGTCAATCTA
GTCGGAGTACTGTCCTGTTTTAGAGCTAGAAATAGCAAGTTAAAATAAGGCTAGTCCGTTAT
CAACTTGAAAAAGTGGCACCGAGTCGGTGC

Modular iSBH-sgRNA (trigger C/sgRNA2/loop=18nt):
GCAGGACAGTACAGCGAACAGACAGACTGTGAAAGGAGAGGCGAGGGAGGTCAATGTCAATC
TAGTCGGAGTACTGTCCTGTTTTAGAGCTAGAAATAGCAAGTTAAAATAAGGCTAGTCCGTT
ATCAACTTGAAAAAGTGGCACCGAGTCGGTGC

Modular iSBH-sgRNA (trigger C/sgRNA2/loop=20nt):
GCAGGACAGTACAGCGAACAGACAGACTGTGAAAGGAGAGGCGAGGGAGGAGTCAATGTCGT
TCTAGTCGGAGTACTGTCCTGTTTTAGAGCTAGAAATAGCAAGTTAAAATAAGGCTAGTCCG
TTATCAACTTGAAAAAGTGGCACCGAGTCGGTGC

Modular iSBH-sgRNA (trigger C/sgRNA2/loop=22nt):
GCAGGACAGTACAGCGAACAGACAGACTGTGAAAGGAGAGGCGAGGGAGGAGGGTCAATGTC
AATCTAGTCGGAGTACTGTCCTGTTTTAGAGCTAGAAATAGCAAGTTAAAATAAGGCTAGTC
CGTTATCAACTTGAAAAAGTGGCACCGAGTCGGTGC

Modular iSBH-sgRNA (trigger C/sgRNA2/loop=24nt):
GCAGGACAGTACAGCGAAGACAGACTGTGAAAGGAGAGGCGAGGGAGGAGGGATGATTTATC
AGGATGTAGTCGGAGTACTGTCCTGTTTTAGAGCTAGAAATAGCAAGTTAAAATAAGGCTAG
TCCGTTATCAACTTGAAAAAGTGGCACCGAGTCGGTGC

Modular iSBH-sgRNA (trigger C/sgRNA2/loop=26nt):
GCAGGACAGTACAGCGAACAGACAGACTGTGAAAGGAGAGGCGAGGGAGGAGGGATGATCAA
TGTCGTTCTAGTCGGAGTACTGTCCTGTTTTAGAGCTAGAAATAGCAAGTTAAAATAAGGCT
AGTCCGTTATCAACTTGAAAAAGTGGCACCGAGTCGGTGC

Modular iSBH-sgRNA (trigger C/sgRNA2/loop=32nt):
GCAGGACAGTACAGCGAGACTTTGACAGACAGACTGTGAAAGGAGAGGCGAGGGAGGAGGGA
TGCTGGATGTATAAGAGTCGGAGTACTGTCCTGTTTTAGAGCTAGAAATAGCAAGTTAAAAT
AAGGCTAGTCCGTTATCAACTTGAAAAAGTGGCACCGAGTCGGTGC

Modular iSBH-sgRNA (trigger D/sgRNA2/loop=14nt (1)):
GCAGGACAGTACAGCGAGACAGTGTTCTCTGCACAGATAAGGACAAGCAATGAAATCTGAGT
CGGAGTACTGTCCTGTTTTAGAGCTAGAAATAGCAAGTTAAAATAAGGCTAGTCCGTTATCA
ACTTGAAAAAGTGGCACCGAGTCGGTGC

Modular iSBH-sgRNA (trigger D/sgRNA2/loop=14nt (2)):

GCAGGACAGTACAGCGAAGTGTTCTCTGCACAGATAAGGACAAACATGTTAAGACTACAAGT
CGGAGTACTGTCCTGTTTTAGAGCTAGAAATAGCAAGTTAAAATAAGGCTAGTCCGTTATCA
ACTTGAAAAAGTGGCACCGAGTCGGTGC

Modular iSBH-sgRNA (trigger D/sgRNA2/loop=16nt):
GCAGGACAGTACAGCGAGACAGTGTTCTCTGCACAGATAAGGACAAACGCAATGAAATCTGA
GTCGGAGTACTGTCCTGTTTTAGAGCTAGAAATAGCAAGTTAAAATAAGGCTAGTCCGTTAT
CAACTTGAAAAAGTGGCACCGAGTCGGTGC

Modular iSBH-sgRNA (trigger D/sgRNA2/loop=18nt):
GCAGGACAGTACAGCGAAGACAGTGTTCTCTGCACAGATAAGGACAAACACAGTTAACTATG
TAGTCGGAGTACTGTCCTGTTTTAGAGCTAGAAATAGCAAGTTAAAATAAGGCTAGTCCGTT
ATCAACTTGAAAAAGTGGCACCGAGTCGGTGC

Modular iSBH-sgRNA (trigger D/sgRNA2/loop=20nt):
GCAGGACAGTACAGCGAAGCAAATGTGTTGCCAAAAAGGATGCTTTAGAGACGGCTTCACTG
TTGAGTCGGAGTACTGTCCTGTTTTAGAGCTAGAAATAGCAAGTTAAAATAAGGCTAGTCCG
TTATCAACTTGAAAAAGTGGCACCGAGTCGGTGC

Modular iSBH-sgRNA (trigger D/sgRNA2/loop=22nt):
GCAGGACAGTACAGCGAGAGACAGTGTTCTCTGCACAGATAAGGACAAACATTAAGATTACA
AGGTCAGTCGGAGTACTGTCCTGTTTTAGAGCTAGAAATAGCAAGTTAAAATAAGGCTAGTC
CGTTATCAACTTGAAAAAGTGGCACCGAGTCGGTGC

Modular iSBH-sgRNA (trigger D/sgRNA2/loop=24nt):
GCAGGACAGTACAGCGAAGAGACAGTGTTCTCTGCACAGATAAGGACAAACATTATGAGTTC
ACGATCTAGTCGGAGTACTGTCCTGTTTTAGAGCTAGAAATAGCAAGTTAAAATAAGGCTAG
TCCGTTATCAACTTGAAAAAGTGGCACCGAGTCGGTGC

Modular iSBH-sgRNA (trigger D/sgRNA2/loop=34nt):
GCAGGACAGTACAGCGAGACAGTGTTCTCTGCACAGATAAGGACAAACATTATTCAGAGGGA
GTACGCATTGAAATCTGAGTCGGAGTACTGTCCTGTTTTAGAGCTAGAAATAGCAAGTTAAA
ATAAGGCTAGTCCGTTATCAACTTGAAAAAGTGGCACCGAGTCGGTGC

*An additional GC sequence was added to all iSBH-sgRNA sequences. G helps expression from U6 promoter, while C base-pairs with the first C from the scaffold

#### 6. RNA triggers for first-generation iSBH-sgRNAs

hairpin linker 34nt trigger 100nt 3’ flank linker hairpin

First-generation iSBH-sgRNA 1 trigger:
GGCTCGAAAGAGCCACGGATGCGTAGTGTGCAGAAGCGACTAGCGAAGCAATGCTAGTACGA
AAGTACTAGC

First-generation iSBH-sgRNA 2 trigger:
GGCTCGAAAGAGCCACTGTTAGGGATGTTCGTATCTCGCTGTACTGTCCTGCATGCTAGTAC
GAAAGTACTAGC

*GC complementary with extra GC sequences on the iSBH-sgRNA 1

First-generation iSBH-sgRNA 3 trigger:
GGCTCGAAAGAGCCACTGTATTCACTGGTCCATTCCATACTTCCCGCCTCATGCTAGTACGA
AAGTACTAGC

First-generation iSBH-sgRNA 4 trigger:
GGCTCGAAAGAGCCACTTGATGTGGGGAGCCCATTTAAGTACTAGCAGGAATGCTAGTACGA
AAGTACTAGC

First-generation iSBH-sgRNA 5 trigger:
GGCTCGAAAGAGCCACTGGACGAACAAGATGCAGAACTGGAGTAAGAGTCATGCTAGTACGA
AAGTACTAGC

First-generation iSBH-sgRNA 1 trigger 100nt 3’ flank:
GGCTCGAAAGAGCCACTGAGACGTAAGACTCGAGCGGTCCGTCTCAGTCAGTATGGTCTTGG
GATGCGTAGTGTGCAGAAGCGACTAGCGAAGCAGGCGGCCGTATGAGGGACAATTGGAGaag
tgaattatataaatataaagtagtaaaaattgaaccattaggagtagcacccaccaagGCAA
AGAGAAGAGTGGTGCAGGTACGAAAGTAC

First-generation iSBH-sgRNA 2 trigger 100nt 3’ flank:
GGCTCGAAAGAGCCACTGAGACGTAAGACTCGAGCGGTCCGTCTCAGTCATAGTATATGTGT
GTTAGGGATGTTCGTATCTCGCTGTACTGTCCTGGCGGCCGCATAGTTAAGCCAGTATCTGC
tccctgcttgtgtgttggaggtcgctgagtagtgcgcgagcaaaatttaagctacaacaaGG
CAAGGCTTGACCGACAAGTACGAAAGTAC

First-generation iSBH-sgRNA 3 trigger 100nt 3’ flank:
GGCTCGAAAGAGCCACTGAGACGTAAGACTCGAGCGGTCCGTCTCAGTCACTATCTATATCT
GTATTCACTGGTCCATTCCATACTTCCCGCCTCGGCGGCCGTCGGGAGATCTCCCGATCCCC
tatggtgcactctcagtacaatctgctctgatgccgcatagttaagccagtatctGCTCCCT
GCTTGTGTGTTGGAGGTGTACGAAAGTAC

#### 7. RNA triggers for second-generation iSBH-sgRNAs

hairpin linker 100nt 5’ flank 44nt trigger 100nt 3’ flank
linker hairpin

Second-generation iSBH-sgRNA 1 trigger:
GGCTCGAAAGAGCCACGGATGCGTAGTGTGAACGTACGGACAGAAGCGACTAGCGAAGCAAT
GCTAGTACGAAAGTACTAGC

Second-generation iSBH-sgRNA 2 trigger:
GGCTCGAAAGAGCCACTGTTAGGGATGTTCGACTAAGCTGGTATCTCGCTGTACTGTCCTAT
GCTAGTACGAAAGTACTAGC

Second-generation iSBH-sgRNA 3 trigger:
GGCTCGAAAGAGCCACTGTATTCACTGGTCCCATCCCACTCATTCCATACTTCCCGCCTCAT
GCTAGTACGAAAGTACTAGC

Second-generation iSBH-sgRNA 1 trigger 100nt 3′ flank:
GGCTCGAAAGAGCCACTGAGACGTAAGACTCGAGCGGTCCGTCTCAGTCAGTATGGTCTTGG
GATGCGTAGTGTGAACGTACGGACAGAAGCGACTAGCGAAGCAGCGGCCGTATGAGGGACAA
TTGGAGaagtgaattatataaatataaagtagtaaaaattgaaccattaggagtagcaccca
ccaagGCAAAGAGAAGAGTGGTGCAGGTACGAAAGTAC

Second-generation iSBH-sgRNA 2 trigger 100nt 3′ flank:
GGCTCGAAAGAGCCACTGAGACGTAAGACTCGAGCGGTCCGTCTCAGTCATAGTATATGTGT
GTTAGGGATGTTCGACTAAGCTGGTATCTCGCTGTACTGTCCTGCGGCCGCATAGTTAAGCC
AGTATCTGCtccctgcttgtgtgttggaggtcgctgagtagtgcgcgagcaaaatttaagct
acaacaaGGCAAGGCTTGACCGACAAGTACGAAAGTAC

Second-generation iSBH-sgRNA 3 trigger 100nt 3′ flank:
GGCTCGAAAGAGCCACTGAGACGTAAGACTCGAGCGGTCCGTCTCAGTCATCGGTCTTGTCT
GTATTCACTGGTCCCATCCCACTCATTCCATACTTCCCGCCTCGCGGCCGTCGGGAGATCTC
CCGATCCCCtatggtgcactctcagtacaatctgctctgatgccgcatagttaagccagtat
ctGCTCCCTGCTTGTGTGTTGGAGGTGTACGAAAGTAC

Second-generation iSBH-sgRNA 1 trigger 100nt 5′ flank:
GGCTCGAAAGAGCCACCCTATGGTGCACTCTCAGTACaatctgctctgatgccgcatagtta
agccagtatctgctccctgcttgtgtgttggaGGTCGCTGAGTAGTGCGCGAGCGTCAGTAT
GGTCTTGGGATGCGTAGTGTGAACGTACGGACAGAAGCGACTAGCGAAGCAGCGGCCGCATG
CTAGTACGAAAGTACTAGC

Second-generation iSBH-sgRNA 2 trigger 100nt 5′ flank:
GGCTCGAAAGAGCCACGCAAAATTTAAGCTACAACAAGGCaaggcttgaccgacaattgcat
gaagaatctgcttagggttaggcgttttgcgCTGCTTCGCGATGTACGGGCCAGGTCATAGT
ATATGTGTGTTAGGGATGTTCGACTAAGCTGGTATCTCGCTGTACTGTCCTGCGGCCGCATG
CTAGTACGAAAGTACTAGC

Second-generation iSBH-sgRNA 3 trigger 100nt 5′ flank:
GGCTCGAAAGAGCCACGTCGACGGATCGGGAGATCTCCCGATCCCctatggtgcactctcag
tacaatctgctctgatgccgcatagttaagcCAGTATCTGCTCCCTGCTTGTGTGTCATCGG
TCTTGTCTGTATTCACTGGTCCCATCCCACTCATTCCATACTTCCCGCCTCGCGGCCGCATG
CTAGTACGAAAGTACTAGC

Second-generation iSBH-sgRNA 1 trigger 100nt 5′+3′ flank:
GGCTCGAAAGAGCCACCCTATGGTGCACTCTCAGTACAATCTGCTCTGATGCCGCATAGTTA
AGCCAGTATCTGCTCCCTGCTTGTGTGTTGGAGGTCGCTGAGTAGTGCGCGAGCGTCAGTAT
GGTCTTGGGATGCGTAGTGTGAACGTACGGACAGAAGCGACTAGCGAAGCAGCGGCCGTATG
AGGGACAATTGGAGAAGTGAATTATATAAATATAAAGTAGTAAAAATTGAACCATTAGGAGT
AGCACCCACCAAGGCAAAGAGAAGAGTGGTGCAGGTACGAAAGTAC

Second-generation iSBH-sgRNA 2 trigger 100nt 5′+3′ flank:
GGCTCGAAAGAGCCACGCAAAATTTAAGCTACAACAAGGCAAGGCTTGACCGACAATTGCAT
GAAGAATCTGCTTAGGGTTAGGCGTTTTGCGCTGCTTCGCGATGTACGGGCCAGGTCATAGT
ATATGTGTGTTAGGGATGTTCGACTAAGCTGGTATCTCGCTGTACTGTCCTGCGGCCGCATA
GTTAAGCCAGTATCTGCTCCCTGCTTGTGTGTTGGAGGTCGCTGAGTAGTGCGCGAGCAAAA
TTTAAGCTACAACAAGGCAAGGCTTGACCGACAAGTACGAAAGTAC

Second-generation iSBH-sgRNA 3 trigger 100nt 5′+3′ flank:

GGCTCGAAAGAGCCACGTCGACGGATCGGGAGATCTCCCGATCCCCTATGGTGCACTCTCAG
TACAATCTGCTCTGATGCCGCATAGTTAAGCCAGTATCTGCTCCCTGCTTGTGTGTCATCGG
TCTTGTCTGTATTCACTGGTCCCATCCCACTCATTCCATACTTCCCGCCTCGCGGCCGTCGG
GAGATCTCCCGATCCCCTATGGTGCACTCTCAGTACAATCTGCTCTGATGCCGCATAGTTAA
GCCAGTATCTGCTCCCTGCTTGTGTGTTGGAGGTGTACGAAAGTAC

#### 8. Truncated RNA triggers for second-generation iSBH-sgRNAs

hairpin linker trigger randomized nt linker hairpin
trigger + randomized nt= 44nt

Second-generation iSBH-sgRNA 1 trigger (D1):
GGCTCGAAAGAGCCACGGATGCGTAGTGTGAACGTACGGAATCCGTGATACTAACGCCGGAT
GCTAGTACGAAAGTACTAGC

Second-generation iSBH-sgRNA 1 trigger (D2):
GGCTCGAAAGAGCCACGGATGCGTAGTGTGAACGTACGGACAGAATGATACTAACGCCGGAT
GCTAGTACGAAAGTACTAGC

Second-generation iSBH-sgRNA 2 trigger (D2):
GGCTCGAAAGAGCCACTGTTAGGGATGTTCGACTAAGCTGACGCGAATTGCGCTGCAATAAT
GCTAGTACGAAAGTACTAGC

Second-generation iSBH-sgRNA 2 trigger (D1):
GGCTCGAAAGAGCCACTGTTAGGGATGTTCGACTAAGCTGGTATCAATTGCGCTGCAATAAT
GCTAGTACGAAAGTACTAGC

Second-generation iSBH-sgRNA 3 trigger (D1):
GGCTCGAAAGAGCCACTGTATTCACTGGTCCCATCCCACTATGAATCGCTGAGATAGGAAAT
GCTAGTACGAAAGTACTAGC

Second-generation iSBH-sgRNA 4 trigger (D2):
GGCTCGAAAGAGCCACTGTATTCACTGGTCCCATCCCACTCATTCTCGCTGAGATAGGAAAT
GCTAGTACGAAAGTACTAGC

#### 9. Endogenous trigger sequences for modular iSBH-sgRNA designs

hairpin linker endogenous trigger sequence linker hairpin

RNA trigger A:
GGCTCGAAAGAGCCACCTGATGATCAGTTTAGTCGTGTTGTGGTGATGATCAGTTTAGTTGT
GTTGTGGTGATGATCAGTTTAGTTGTGTTGTGGTGATGATCAGTTTAGTCGTGTTGTGGTGA
TGATCAGTTTAGTCGTGTTGTGGTGATGATCTGTTTAGATGCTAGTACGAAAGTACTAGC

RNA trigger B:
GGCTCGAAAGAGCCACCTCAACAAAAGGTTAAAAATGGTTTAGTCACATGTATGCTAACTTT
CAATCTACAAAACTACAACCATCATCATAATCCCACAATCCTTCCTTCCTCTGTTTCACTTC
ATCTGCCCCCATCAAAGAAAGCAGGGGATGACTAGAGACATTTGGTCGTTGTGATCGCCCGC

CGTGAAATTGGCCCTTGGGCACGGATGATCGGACGTAAGTCTCAAACGCCGGTTGGTGAGCT
GTGAGGTGTTGGATTTCTGCGTTTATTACTGATAATGCTAGTACGAAAGTACTAGC

RNA trigger C:
GGCTCGAAAGAGCCACTCTTAAAGTGCCAGCTTTCCCCTTGAAGTTCTGCCTAAGGTCATCC
CTCCTCCCTCGCCTCTCCTTTCACAGTCTGTCTGTCAAAGTCCCGTCAGTCAGCCCGAGCCC
GCGCCCCCGCCCCTGCACGCGCCTCGGGCACTCGCGTTCTGTCGCTCTGGAATGAGCGAGCG
AGCGAGAGAGAGAGGCGCGCTGCGCACTCCCGACTGGACAACAAGACTGTTTGTGGTTCTGA
GACGCGAAAGCGAATGTGGGTCAAAATCTGACAAGATGCTAGTACGAAAGTACTAGC

RNA trigger D:
GGCTCGAAAGAGCCACCACTGGCTTAGGAGTCTCAGCTCTGGGTACTCCCTCTGAATAATGT
TTGTCCTTATCTGTGCAGAGAACACTGTCTCTAAAGCATCCTTTGGCAACACATTTGCTCAA
TCAACTACTGAATTGGTTGTTAAAATTAATTTTCCTTTATGCTAGTACGAAAGTACTAGC

#### 10. 1xCTS sequences

**sequence matching sgRNA PAM site**

1xCTS sgRNA 1: CAGTCGCGTGTAGCGAAGCAAGG

1xCTS sgRNA 2: GTAAGTCGGAGTACTGTCCTAGG

1xCTS sgRNA 3: CATGACATTATTCCCGCCTCAGG

1xCTS sgRNA 4: CCAGGTAATGACTAGCAGGAAGG

1xCTS sgRNA 5: GCACTACTTCAGTAAGAGTCAGG

#### 11. 8xCTS sequences

**Linker 8x(sequence matching sgRNA PAM site linker) linker**

8xCTS sgRNA 1:
TCTAGAGAGCTCCAGTCGCGTGTAGCGAAGCACGGCTATGAAAATATTAACAGTCGCGTGTA
GCGAAGCAGGGACTAGTAGGGTCAGTAGGAGCTCCAGTCGCGTGTAGCGAAGCACGGCTATG
AAAATATTAACAGTCGCGTGTAGCGAAGCAGGGACTAGTGGGCCCCTAGAGAGCTCCAGTCG
CGTGTAGCGAAGCACGGCTATGAAAATATTAACAGTCGCGTGTAGCGAAGCAGGGACTAGTA
GGGTCAGTAGGAGCTCCAGTCGCGTGTAGCGAAGCACGGCTATGAAAATATTAACAGTCGCG
TGTAGCGAAGCAGGGACTAGTGCTAGC

8xCTS sgRNA 2:
TCTAGAGAGCTCGTAAGTCGGAGTACTGTCCTCGGCTATGAAAATATTAAGTAAGTCGGAGT
ACTGTCCTGGGACTAGTAGGGTCAGTAGGAGCTCGTAAGTCGGAGTACTGTCCTCGGCTATG
AAAATATTAAGTAAGTCGGAGTACTGTCCTGGGACTAGTGGGCCCCTAGAGAGCTCGTAAGT
CGGAGTACTGTCCTCGGCTATGAAAATATTAAGTAAGTCGGAGTACTGTCCTGGGACTAGTA
GGGTCAGTAGGAGCTCGTAAGTCGGAGTACTGTCCTCGGCTATGAAAATATTAAGTAAGTCG
GAGTACTGTCCTGGGACTAGTGCTAGC

8xCTS sgRNA 3:
TCTAGAGAGCTCCATGACATTATTCCCGCCTCCGGCTATGAAAATATTAACATGACATTATT
CCCGCCTCGGGACTAGTAGGGTCAGTAGGAGCTCCATGACATTATTCCCGCCTCCGGCTATG
AAAATATTAACATGACATTATTCCCGCCTCGGGACTAGTGGGCCCCTAGAGAGCTCCATGAC
ATTATTCCCGCCTCCGGCTATGAAAATATTAACATGACATTATTCCCGCCTCGGGACTAGTA
GGGTCAGTAGGAGCTCCATGACATTATTCCCGCCTCCGGCTATGAAAATATTAACATGACAT
TATTCCCGCCTCGGGACTAGTGCTAG

8xCTS sgRNA 4:
TCTAGAGAGCTCCCAGGTAATGACTAGCAGGACGGCTATGAAAATATTAACCAGGTAATGAC
TAGCAGGAGGGACTAGTAGGGTCAGTAGGAGCTCCCAGGTAATGACTAGCAGGACGGCTATG
AAAATATTAACCAGGTAATGACTAGCAGGAGGGACTAGTGGGCCCCTAGAGAGCTCCCAGGT
AATGACTAGCAGGACGGCTATGAAAATATTAACCAGGTAATGACTAGCAGGAGGGACTAGTA
GGGTCAGTAGGAGCTCCCAGGTAATGACTAGCAGGACGGCTATGAAAATATTAACCAGGTAA
TGACTAGCAGGAGGGACTAGTGCTAG

8xCTS sgRNA 5:
TCTAGAGAGCTCGCACTACTTCAGTAAGAGTCCGGCTATGAAAATATTAAGCACTACTTCAG
TAAGAGTCGGGACTAGTAGGGTCAGTAGGAGCTCGCACTACTTCAGTAAGAGTCCGGCTATG
AAAATATTAAGCACTACTTCAGTAAGAGTCGGGACTAGTGGGCCCCTAGAGAGCTCGCACTA
CTTCAGTAAGAGTCCGGCTATGAAAATATTAAGCACTACTTCAGTAAGAGTCGGGACTAGTA
GGGTCAGTAGGAGCTCGCACTACTTCAGTAAGAGTCCGGCTATGAAAATATTAAGCACTACT
TCAGTAAGAGTCGGGACTAGTGCTAG

*8xCTS sequences displayed here are in reverse orientation

#### 12. Chemically modified sgRNAs and iSBH-sgRNAs tested in zebrafish

GC spacer* extension* loop extension spacer scaffold extra Us
Backfold=spacer*+extension+loop

*An additional GC sequence was added to all iSBH-sgRNA sequences to maintain consistency.

sgRNA 2 with chemical modifications
mG*mU*mA*rArGrUrCrGrGrArGrUrArCrUrGrUrCrCrUrGrUrUrUrArGrArGrCrUrAr
GrArArArUrArGrCrArArGrUrUrArArArArUrArArGrGrCrUrArGrUrCrCrGrUrUmAmUm
CmAmAmCmUmUmGmAmAmAmAmGmUmGmGmCmAmCmCmGmAmGmUm
CmGmGmUmGmCmU*mU*mU*rU

Second-generation iSBH-sgRNA 2 with chemical modifications (first iteration):
mG*mC*mA*rGrGrArCrArGrUrArCrArGrCrGrArGrArUrArCrCrArGrCrUrUrArGrUrCrG
rArArCrArUrCrCrCrUrArArCrArGrArCrArUrArGrCrCrCrGrUrArArGrUrCrGrGrArGrUr
ArCrUrGrUrCrCrUrGrUrUrUrUrArGrArGrCrUrArGrArArArUrArGrCrArArGrUrUrArArA
rArUrArArGrGrCrUrArGrUrCrCrGrUrUmAmUmCmAmAmCmUmUmGmAmAmAmA
mAmGmUmGmGmCmAmCmCmGmAmGmUmCmGmGmUmGmCmU*mU*mU*rU

Second-generation iSBH-sgRNA 2 with chemical modifications (second iteration):
mG*mC*mA*rGrGrArCrArGrUrArCrArGrCrGrArGrArUrArCrCrArGrCrUrUrArGrUrCrG
rArArCrArUrCrCrCrUrArArCrArGrArCrArUrArGrCrCrCmG*mU*mA*rArGrUrCrGrGrAr
GrUrArCrUrGrUrCrCrUrGrUrUrUrArGrArGrCrUrArGrArArArUrArGrCrArArGrUrUrA

rArArArUrArArGrGrCrUrArGrUrCrCrGrUrUmAmUmCmAmAmCmUmUmGmAmAmA
mAmAmGmUmGmGmCmAmCmCmGmAmGmUmCmGmGmUmGmCmU*mU*mU*rU

m= 2'OMe RNA base
r= ribobase
*= phosphorothioate bond

#### 13. Chemically modified RNA triggers tested in zebrafish

hairpin linker 34nt trigger 100nt 3' flank linker hairpin
random sequence

Non-complementary RNA trigger with chemical modifications:
mG*mG*mC*rUrCrGrArArArGrArGrCrCrArCrGrGrGrUrCrUrUrCrGrArGrCrArUrUrGr
GrArArGrUrCrUrArGrArUrArCrCrUrGrCrUrUrUrArCrGrArArGrArCrCrUrArUrGrCrUrA
rGrUrArCrGrArArArGrUrArCrU*mA*mG*mC

Second-generation iSBH-sgRNA 2 trigger with chemical modifications:
mG*mG*mC*rUrCrGrArArArGrArGrCrCrArCrUrGrUrUrArGrGrGrArUrGrUrUrCrGrAr
CrUrArArGrCrUrGrGrUrArUrCrUrCrGrCrUrGrUrArCrUrGrUrCrCrUrArUrGrCrUrArGr
UrArCrGrArArArGrUrArCrU*mA*mG*mC

m= 2'OMe RNA base
r= ribobase
*= phosphorothioate bond

#### 14. Oligos and primer sequences ordered for cloning 8xCTS reporters

Oligo 2xCTS sgRNA 1:
GGACCCTACTAGTCCCTGCTTCGCTACACGCGACTGTTAATATTTTCATAGCCGTGCTTCGC
TACACGCGACTGGAGCTCCTACTGACCC

Oligo 2xCTS sgRNA 2:
GGACCCTACTAGTCCCAGGACAGTACTCCGACTTACTTAATATTTTCATAGCCGAGGACAGT
ACTCCGACTTACGAGCTCCTACTGACCC

Oligo 2xCTS sgRNA 3:
GGACCCTACTAGTCCCGAGGCGGGAATAATGTCATGTTAATATTTTCATAGCCGGAGGCGGG
AATAATGTCATGGAGCTCCTACTGACCC

Oligo 2xCTS sgRNA 4:
GGACCCTACTAGTCCCTCCTGCTAGTCATTACCTGGTTAATATTTTCATAGCCGTCCTGCTA
GTCATTACCTGGGAGCTCCTACTGACCC

Oligo 2xCTS sgRNA 5:
GGACCCTACTAGTCCCGACTCTTACTGAAGTAGTGCTTAATATTTTCATAGCCGGACTCTTA
CTGAAGTAGTGCGAGCTCCTACTGACCC

Primers used for amplifying 2xCTS oligos and cloning 4xCTS-ECFP reporters

OP_2xCTS_FW: ACTGCCGTCTCGGACCCTACTAGTCCC
OP_2xCTS_RW: ACCTACGTCTCGGGTCAGTAGGAGCTC

Primers used for amplifying 4xCTS plasmids and cloning 8xCTS-ECFP reporters
OP_4xCTS_NheI_FW: AGAGATACCGGGCGCTAGCACTAGTCCC
OP_4xCTS_ApaI_RW: GCCCCCCGCTAGTGGGCCCCTAGAGAGCTC

#### 15. Oligos and primer sequences ordered for repetitive trigger sequences

**Trigger A**

**OP_trig_A_oligo_1:**
**GAAGAGGAAGACCTCCACCTGATGATCAGTTTAGTCGTGTTGTGGTGATGATCAGTTTAGTT GTGTTGCTGTCTTCGCACTACGC**

OP_trig_A_oligo_1_FW_BbsI: GAAGAGGAAGACCTCCAC
OP_trig_A_oligo_1+2_RW_BbsI: GCGTAGTGCGAAGACAG

**OP_trig_A_oligo_2:**
**CCACGACTGGAAGACTGGTTGTGGTGATGATCAGTTTAGTTGTGTTGTGGTGATGATCAGTT TAGTCCTGTCTTCGCACTACGC**

OP_trig_A_oligo_2+3_FW_BbsI: CCACGACTGGAAGACTG

OP_trig_A_oligo_1+2_RW_BbsI: GCGTAGTGCGAAGACAG

**OP_trig_A_oligo_3:**
**CCACGACTGGAAGACTGAGTCGTGTTGTGGTGATGATCAGTTTAGTCGTGTTGTGGTGATGA TCTGTTTAGATGCCGGTCTTCTCGTC**

OP_trig_A_oligo_2+3_FW_BbsI: CCACGACTGGAAGACTG
OP_trig_A_oligo_3_RW_BbsI: GACGAGAAGACCGGCAT

**Trigger B**

**OP_trig_B_oligo_1:**
**CATGTATGCTAACTTTCAATCTACAAAACTACAACCATCATCATAATCCCACAATCCTTCCT TCCTCTGTTTTTCACTTCATCTGCCCCC**

OP_trig_B_oligo_1_FW_BbsI:
AAGAGGAAGACCTCCACCTCAACAAAAGGTTAAAAATGGTTTAGTCACATGTATGCTAACTT TCAATCTAC

OP_trig_B_oligo_1_RW_BbsI:
ATGCAGAAGACCACCTGCTTTCTTTGATGGGGGCAGATGAAGTGAAAAAC

**OP_trig_B_oligo_2:**

**CATTTGGTCGTTGTGATCGCCCGCCGTGAAATTGGCCCTTGGGCACGGATGATCGGACGTAA GTCTCAAACGCCGGTTGGTGAGCTGTGA**

OP_trig_B_oligo_2_FW_BbsI:
ACTGGAAGACTGCAGGGGATGACTAGAGACATTTGGTCGTTGTGATCG

OP_trig_B_oligo_2_RW_BbsI:
TACTAGAAGACTGGCATTATCAGTAATAAACGCAGAAATCCAACACCTCACAGCTCACCAAC
C

#### 16. Primer sequences for RNA circularisation assays

**Second-generation iSBH-sgRNA circularisation assay**

Second-generation iSBH-sgRNA 1 primers

OP_Ca_nes1_F: TTATCAACTTGAAAAAGTGGCAC
OP_Ca_nes1_R+RT: GACTAGCCTTATTTTAACTTGC
(primer also used for RT)
OP_CA_nes2_NheI_F: AAGCTGGCTAGCAAAAGTGGCACCGAGTC
OP_CA_nes2_sgRNA1_NotI_R: ACTCGAGCGGCCGCCTAGCTCTAAAACTGCTTCG

Second-generation iSBH-sgRNA 2 primers

OP_Ca_nes1_F: TTATCAACTTGAAAAAGTGGCAC
OP_Ca_nes1_R+RT: GACTAGCCTTATTTTAACTTGC
(primer also used for RT)
OP_CA_nes2_NheI_F: AAGCTGGCTAGCAAAAGTGGCACCGAGTC
OP_CA_ nes2_sgRNA2_NotI_R: ACTCGAGCGGCCGCCTAGCTCTAAAACAGGACAG

Second-generation iSBH-sgRNA 3 primers

OP_Ca_nes1_F: TTATCAACTTGAAAAAGTGGCAC
OP_Ca_nes1_R+RT: GACTAGCCTTATTTTAACTTGC
(primer also used for RT)
OP_CA_nes2_NheI_F: AAGCTGGCTAGCAAAAGTGGCACCGAGTC
OP_CA_nes2_sgRNA3_NotI_R: ACTCGAGCGGCCGCCTAGCTCTAAAACGAGGCG

**RNA triggers circularisation assay**

Second-generation iSBH-sgRNA 1 trigger 100nt 3' flank primers

OP_CA_nes1_trig1_F: CCACCAAGGCAAAGAGAAG

OP_CA_nes1_trig1_R+RT: CTACTCCTAATGGTTCAATTTTTACTAC
(primer also used for RT)
OP_CA_nes2_trig1_F_NheI: AAGCTGGCTAGCCAAAGAGAAGAGTGGTGCAG
OP_CA_nes2_trig1_R_NotI:
ACTCGAGCGGCCGCCAATTTTTACTACTTTATATTTATATAATTCACTTCTC

Second-generation iSBH-sgRNA 2 trigger 100nt 3' flank primers

OP_CA_nes1_trig2_F: TACAACAAGGCAAGGCTTG
OP_CA_nes1_trig2_R+RT: CTTAAATTTTGCTCGCGCAC
(primer also used for RT)
OP_CA_nes2_trig2_F_NheI: AAGCTGGCTAGCCAAGGCTTGACCGACAAG
OP_CA_nes2_trig2_R_BbsI: GAGCCGAAGACCCGGCCGCCGACCTCCAACACACAAG

Second-generation iSBH-sgRNA 3 trigger 100nt 3' flank primers

OP_CA_nes1_trig3_F: AGTATCTGCTCCCTGCTTG
OP_CA_nes1_trig3_R+RT: TTAACTATGCGGCATCAGAG
(primer also used for RT)
OP_CA_nes2_trig3_F_NheI: AAGCTGGCTAGCGCTTGTGTGTTGGAGGTG
OP_CA_nes2_trig3_R_NotI: ACTCGAGCGGCCGCGAGCAGATTGTACTGAGAGTG

#### 17. Primer sequences- zebrafish genotyping

OP_dCas9_Vp64_nes1_F: AAATAAGCGAATTCTCCAAAAG
OP_dCas9_Vp64_nes1_R:CTACGTCTCGCCGCATGTTAGCAGACTTCCTCTGCCCTCCG
AGCCACCGCCCTTGTACAGCTCGTCCATG
OP_dCas9_Vp64_nes2_F: TGCTTGGTTCGGATGCCCTTGATGAC
OP_dCas9_Vp64_nes2_R: GGCGAAGCACATCAGGCCGTAGC
OP_8xCTS _nes1_F: GGTACGGGAGGTACTTGGAGCG
OP_8xCTS _nes1_R: CACCACCCCGGTGAACAGCTC
OP_8xCTS _nes2_F: CTTGGAGCGGCCGCAATAAATATC
OP_8xCTS _nes2_R: CTTGCTCACCATGGTCAACGC

#### Appendix 2- Figure Supplements

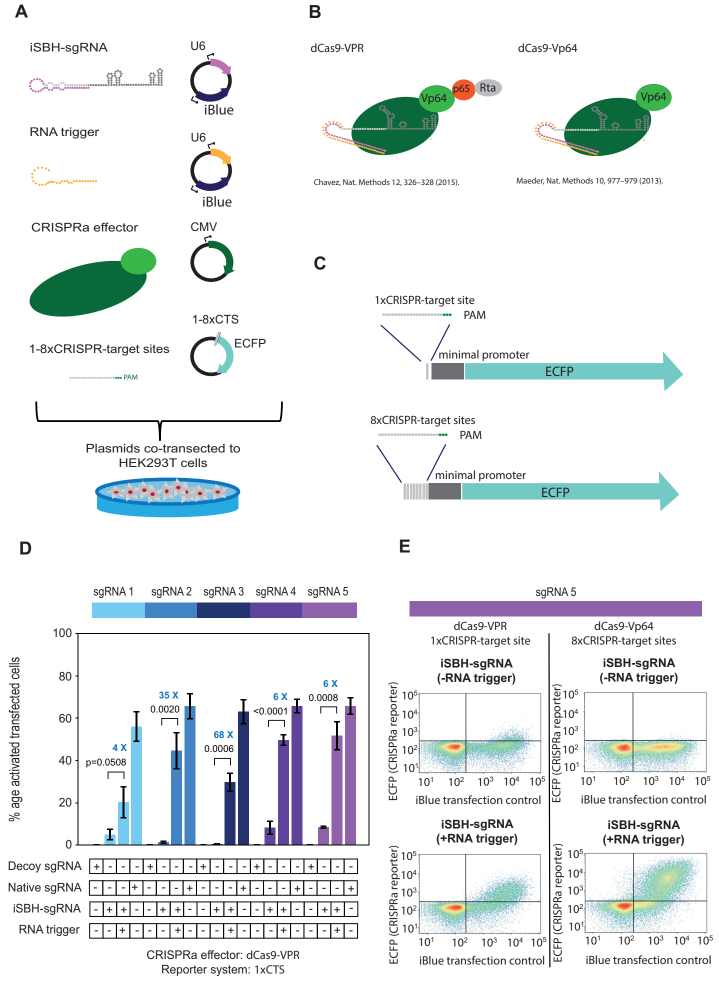


**Figure 1—figure supplement 1.** First generation iSBH-sgRNAs detect short RNA triggers in HEK293T cells. **A.** iSBH-sgRNAs and RNA triggers were cloned in mammalian U6 expression vectors. 1 or 8 CRISPR-target sequences (CTSs) were also cloned upstream of an ECFP fluorescent reporter. We co-transfected the 3 cloned plasmids as well as CRISPRa effectors into HEK293T cells. Fluorescence outputs were measured by Flow Cytometry. **B.** Effectors used include dCas9-VPR (***Chavez et al.*** (***2015***)) and dCas9-Vp64 (***Maeder et al.*** (***2013***)). In contrast with dCas9-Vp64, dCas9-VPR is a stronger transcriptional activator due to the presence of the additional p65 and Rta domains. **C.** 1xCTS (CRISPR-target sequence) reporters contain a single Cas9-binding region upstream from the ECFP reporter. 8xCTS reporters (***Nissim et al.*** (***2014***)) contain 8 repeats of the Cas9-binding region upstream from the ECFP reporter. For promoting expression of the ECFP reporters, minimal adenovirus major late promoters were also introduced within these cassettes. **D.** Starting from five different sgRNA spacer sequences, we designed 5 different iSBH-sgRNA sequences. For each iSBH-sgRNA, corresponding RNA triggers and 1xCTS-ECFP reporters were also designed. Ability of first-generation iSBH-sgRNA designs to drive expression of the ECFP reporter was assessed in the absence or presence of complementary RNA triggers. Experiments were carried out using dCas9-VPR and 1xCTS-ECFP reporters. **E.** Plots compare ECFP fluorescence observed for iSBH-sgRNA 5 activation. In the left, OFF-state and ON-state activity is displayed using CRISPRa assays consisting of the 1xCTS reporters and dCas9-VPR. In the right, OFF-state and ON-state activity is displayed using CRISPRa assays consisting of the 8xCTS reporters and dCas9-Vp64. As iBlue fluorescent gene is encoded in plasmids encoding both iSBH-sgRNAs and RNA triggers, this marker serves as a transfection control. Figure shows mean +/- standard deviation values measured for 3 biological replicates. Values above bars represent fold turn-on values for iSBH-sgRNA activation (blue) and *p*-values (black) determined through unpaired t-tests.

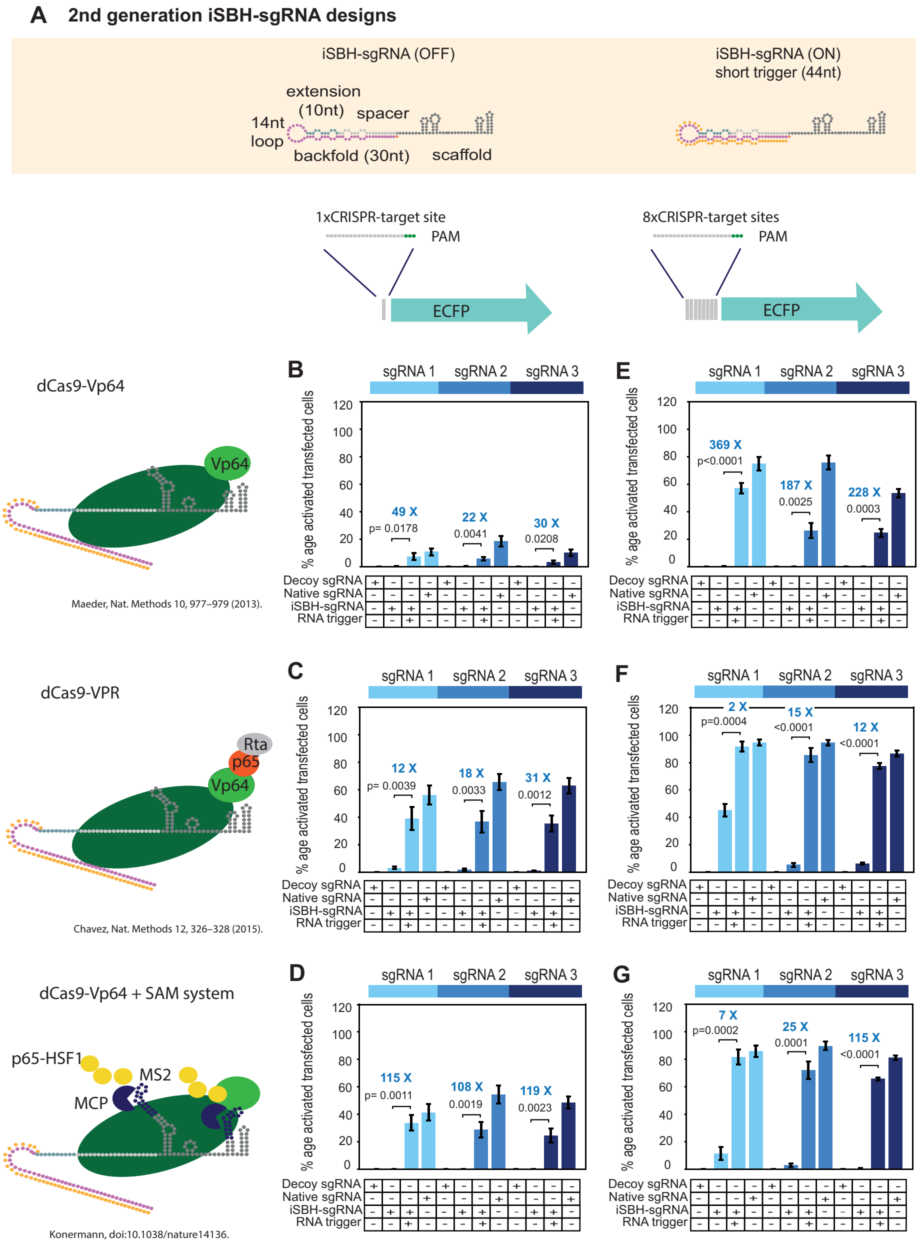


**Figure 2—figure supplement 1.** CRISPRa reporters of choice influence ON/OFF ratios of second generation iSBH-sgRNA designs while detecting short RNA triggers. **A.**Second-generation iSBH-sgRNA designs and corresponding short RNA triggers. **B.**Testing second-generation iSBH-sgRNAs using dCas9-VP64 (***Maeder et al.*** (***2013***)) and 1xCTS-ECFP reporters. **C.**Testing second-generation iSBH-sgRNAs using dCas9-VPR (***Chavez et al.*** (***2015***)) and 1xCTS-ECFP reporters. **D.**Testing second-generation iSBH-sgRNAs using dCas9-VP64, the SAM booster system (***Konermann et al.*** (***2015***)) and 1xCTS-ECFP reporters. **E.**Testing second-generation iSBH-sgRNAs using dCas9-VP64 (***Maeder et al.*** (***2013***)) and 8xCTS-ECFP reporters (***Nissim et al.*** (***2014***)). **F.**Testing second-generation iSBH-sgRNAs using dCas9-VPR (***Chavez et al.*** (***2015***)) and 8xCTS-ECFP reporters (***Nissim et al.*** (***2014***)). **G.**Testing second-generation iSBH-sgRNAs using dCas9-VP64, the SAM booster system (***Konermann et al.*** (***2015***)) and 8xCTS-ECFP reporters (***Nissim et al.*** (***2014***)). Figure shows mean +/- standard deviation values measured for 3 biological replicates. Values above bars represent fold turn-on values for iSBH-sgRNA activation (blue) and *p*-values (black) determined through unpaired t-tests.

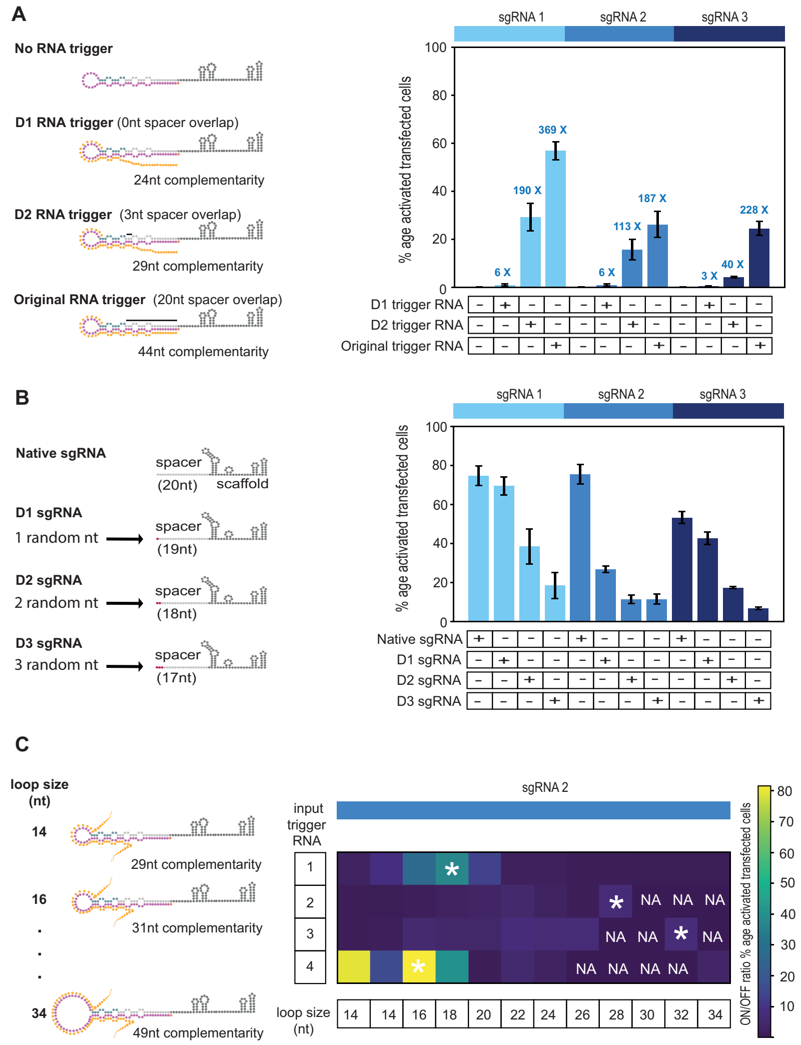


**Figure 3—figure supplement 1.** Modular iSBH-sgRNA designs enable spatial separation of spacer and trigger-sensing sequences. **A.** Trigger RNA truncation experiments. In the original RNA trigger design, the trigger sequence was influenced by the 20nt sgRNA spacer sequence. A way of avoiding this involves to truncating the trigger sequence, so that there is little to no overlap with the spacer sequence. In D1 design, the 20nt spacer sgRNA sequence does not influence at all the sequence of the RNA trigger. In D2 design, the first 3nt of the spacer are still influencing the sequence of the RNA trigger. **B.** Spacer RNA randomisation experiments. Here, we reduced the complementary between the sgRNA spacer sequence and the CTS. This experiment was carried out to accommodate D2 RNA trigger designs and to completely abolish the dependency between the sgRNA spacer and trigger sequences. Native sgRNAs that matched 19, 18 and 17nt of the CTS were designed and tested. **C.** Optimising modular iSBH-sgRNA designs by changing loop length. Multiple MODesign simulations were run for designing iSBH-sgRNAs capable of sensing trigger RNA A (146nt repetitive RNA sequence), trigger RNA B (267nt repetitive RNA sequence), trigger RNA C (268nt repetitive RNA sequence) and trigger RNA D (146nt eRNA sequence). Each simulation contained different loop size specifications and sgRNA 2 sequence was kept constant between simulations. Selected iSBH-sgRNA designs were transfected to HEK293T cells together with corresponding RNA trigger sequences expressed from U6 promoters. For a loop size of 14nt, 2 different modular iSBH-sgRNAs were tested for each trigger. These iSBH-sgRNAs hybridised with different trigger sub-sequences. *NA* represents conditions where the RNA trigger sequence did not allow designing of iSBH-sgRNAs with a particular loop size. iSBH-sgRNA designs with best ON/OFF ratios observed for different RNA triggers are marked with *. Figure shows mean +/- standard deviation values measured for 3 biological replicates.Fold turn-on values for iSBH-sgRNA activation are also displayed in blue.

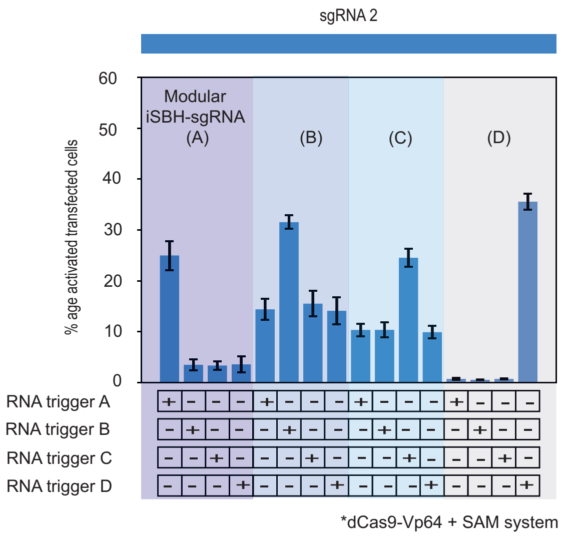


**Figure 3—figure supplement 2.** This supplementary figure reinterprets the data presented in Figure 3.E. using bar plots for enhanced clarity and comparison. It depicts the results of co-transfecting HEK293T cells with four modular iSBH-sgRNAs (A, B, C, and D) and examines all combinations of iSBH-sgRNA: RNA trigger pairings. The bar plots provide a visual representation of mean values with error bars indicating the standard deviation, based on three biological replicates.

891

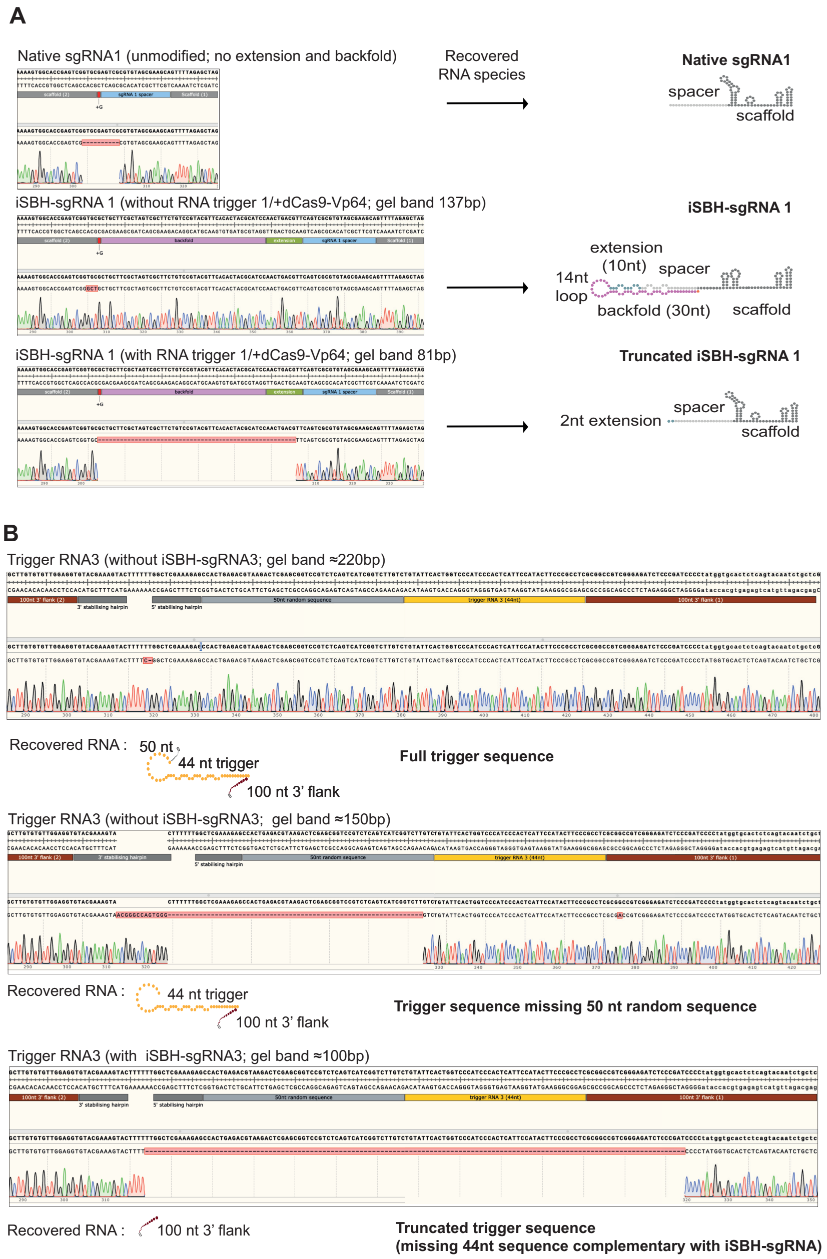


**Figure 4—figure supplement 1.** Sequencing results for iSBH-sgRNA circularisation assays. PCR products were submitted for Sanger Sequencing. Data displays sequencing results obtained for iSBH-sgRNA 1 in the presence of dCas9-Vp64. Top sequencing trace was recovered from cells transfected with non-engineered, native sgRNA. Middle and lower sequencing traces resulted from cells transfected with iSBH-sgRNAs in the absence or presence of complementary RNA triggers. **B.** Sequencing results for RNA trigger circularisation assay. PCR products obtained for RNA trigger 3 were submitted for Sanger Sequencing. Top and middle traces represent RNA trigger sequences recovered in the absence of complementary iSBH-sgRNA (220 and 150bp bands), while bottom trace was recovered in the presence of complementary iSBH-sgRNAs.

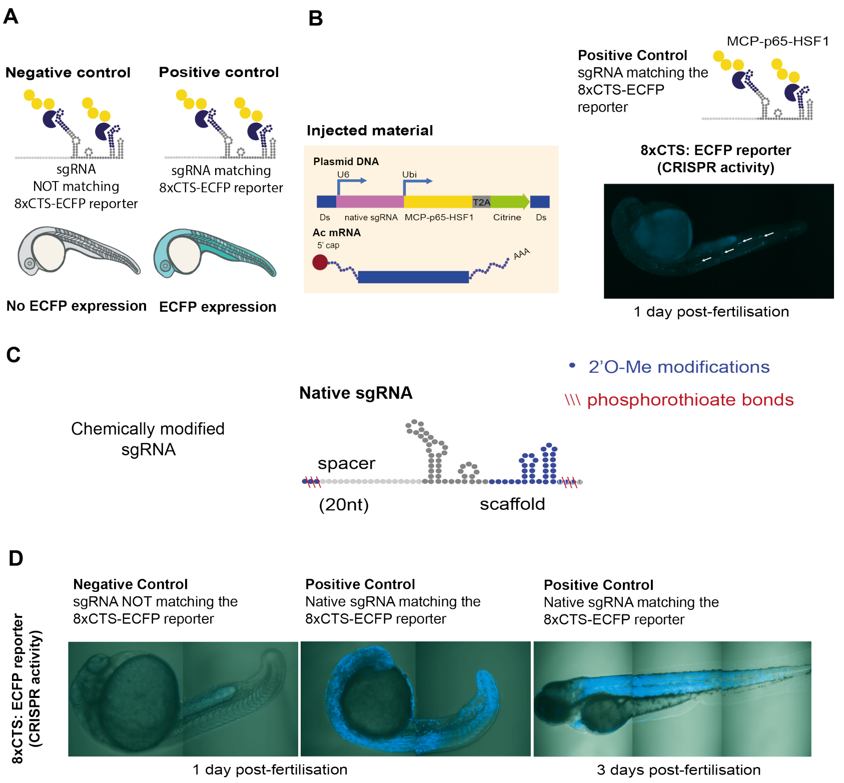


**Figure 5—figure supplement 1.** Optimising sgRNA delivery to zebrafish embryos. **A.** Validation controls. Embryos were injected with native sgRNAs (without the iSBH-sgRNA scaffold) matching or not matching the 8xCTS-ECFP reporter. Only embryos injected with matching sgRNA controls are expected to express ECFP. **B.** Embryos resulted from crossing dCas9-Vp64 and 8xCTS-ECFP lines were injected with control sgRNA sequences. sgRNAs were expressed under the control of a fish U6 promoter and contained the modified sgRNA scaffold compatible with the SAM amplification system. SAM effector proteins (MCP-p63-HSF1, ***Konermann et al.*** (***2015***)) were also encoded into the same vector, together with a Citrine fluorescent protein. This construct was expressed under the control of the ubiquitin promoter. Vectors were co-injected with Ac transposase mRNA (***Chong-Morrison et al.*** (***2018***)). Resulting embryos display mosaic ECFP expression. **C.** sgRNA chemical modifications were designed and synthesised by IDT, comprising 2'O-methyl (Me) modifications as well as phosphorothioate (PS) bonds. **D.** Delivery of chemically modified sgRNAs. Embryos were injected with negative and positive control sgRNAs. For positive control, images are taken at 1 day as well as 3 days post-fertilisation. Resulting embryos display homogeneous ECFP expression across tissues.

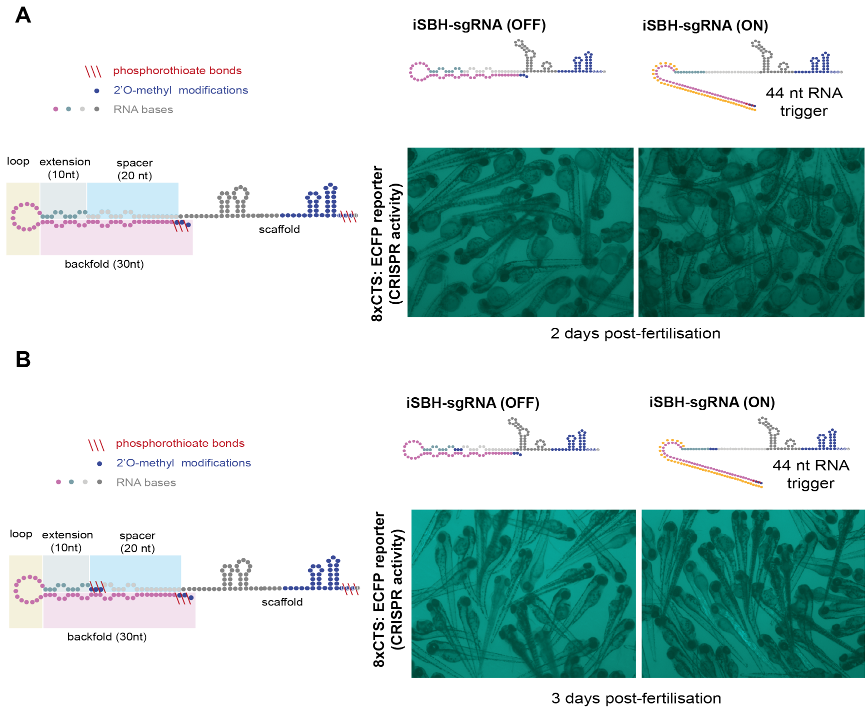


**Figure 5—figure supplement 2.** Testing different iSBH-sgRNA chemical modifications *in vivo.* A. An initial strategy for chemically modifying iSBH-sgRNAs involved protecting the iSBH-sgRNA 5′ end. These modifications were used together with sgRNA scaffold modifications used in the native sgRNA designs (Figure 5- Figure supplement 1.C.). Chemically modified iSBH-sgRNAs were co-injected together with either non-complementary (iSBH-sgRNA OFF) or complementary (iSBH-sgRNA ON) RNA triggers. **B.** A second strategy for chemically modifying iSBH-sgRNAs involved protecting the iSBH-sgRNA 5′ end as well as the 5′end of the sgRNA spacer. These modifications were used together with sgRNA scaffold modifications used in the native sgRNA designs (Figure 5- Figure supplement 1.C.). Chemically modified iSBH-sgRNAs were co-injected together with either non-complementary (iSBH-sgRNA OFF) or complementary (iSBH-sgRNA ON) RNA triggers.
